# Oxygen diffusion in ellipsoidal tumour spheroids

**DOI:** 10.1101/271890

**Authors:** David Robert Grimes, Frederick J. Currell

## Abstract

Oxygen plays a central role in cellular metabolism, in both healthy and tumour tissue. The presence and concentration of molecular oxygen in tumours has a substantial effect on both radiotherapy response and tumour evolution, and as a result the oxygen micro-environment is an area of intense research interest. Multicellular tumour spheroids closely mimic real avascular tumours, and in particular they exhibit physiologically relevant heterogeneous oxygen distribution. This property has made them a vital part of in vitro experimentation. For ideal spheroids, their heterogeneous oxygen distributions can be predicted from theory, allowing determination of cellular oxygen consumption rate (OCR) and anoxic extent. However, experimental tumour spheroids often depart markedly from perfect sphericity. There has been little consideration of this reality. To date, the question of how far an ellipsoid can diverge from perfect sphericity before spherical assumptions breakdown remains unanswered. In this work we derive equations governing oxygen distribution (and more generally, nutrient and drug distribution) in both prolate and oblate tumour ellipsoids, and quantify the theoretical limits of the assumption that the spheroid is a perfect sphere. Results of this analysis yield new methods for quantifying OCR in ellipsoidal spheroids, and how this can be applied to markedly increase experimental throughput and quality.

**Author summary:** Multicellular tumour spheroids (MCTS) are an increasingly important tool in cancer research, exhibiting non-homogeneous oxygen distributions and central necrosis. These are more similar to *in situ* avascular tumours than conventional 2D biology, rendering them exceptionally useful experimental models. Analysis of spheroids can yield vital information about cellular oxygen consumption rates, and the heterogeneous oxygen contribution. However, such analysis pivots on the assumption of perfect sphericity, when in reality spheroids often depart from such an ideal. In this work, we construct a theoretical oxygen diffusion model for ellipsoidal tumour spheroids in both prolate and oblate geometries. With these models established, we quantify the limits of the spherical assumption, and illustrate the effect of this assumption breaking down. Methods of circumventing this breakdown are also presented, and the analysis here suggests new methods for expanding experimental throughput to also include ellipsoidal data.

## Introduction

Oxygen plays a seminal role in cancer treatment and patient prognosis. The presence of molecular oxygen in a tumour markedly increases radio-sensitivity, with well-oxygenated regions responding to radiotherapy up to a factor of three relative to anoxic sub-volumes [1,2]. This oxygen enhancement ratio is also seen in emerging modalities such as proton therapy [3,4], raising the tantalizing prospect of dose-painting, where dose is selectively boosted to hypoxic regions to boost therapy response [5]. The basic idea underpinning dose-painting has been discussed for over a decade, but application has been hampered by difficulty in non-invasive hypoxia imaging. Methods such as F-MISO PET (fluoromisonidazole positron emission tomography) have a maximum resolution in the millimetre regime, while oxygenation varies over a micron scale. As a consequence, mathematical modeling is vital for bridging the resolution gap [6].

Aside from therapeutic considerations, oxygen has a marked impact on patient prognosis. The pioneering work of Gray and colleagues in the early 1950s established that tumour oxygen concentration was correlated with prognosis, and extensive hypoxia was a negative prognostic marker [7]. This finding has been well replicated to present day [8–10] and is not solely due to hypoxia-induced treatment resistance. Under severe hypoxia, tumour cells can respond to such pressure by activating oxygen-sensitive signaling pathways [11,12]. Current biological thinking suggests these signalling pathways act to alter gene expression to promote cell survival under adverse conditions. Hypoxia is also a major driver of angiogenesis, giving rise to new routes for cells to travel along [13,14], endowed with the ability to metastasize [15].

The extraordinary importance of oxygen in cancer treatment and evolution has made it an important avenue of study, with an urgent need for further research. Despite the fundamental importance of molecular oxygen in tumours, investigations have been complicated by the significant experimental difficulty in ascertaining oxygen concentration *in situ* [6]. Real tumours have highly heterogeneous oxygen supply and complex tortured vasculature, and even well-oxygenated regions are frequently inter-spaced with pockets of anoxia [14,16]. Standard 2D monolayers of cells are not an ideal experimental model, typically exhibiting an unrealistically homogeneous oxygen contribution. There is however a more realistic experimental option in the form of tumour spheroids. These clusters of cancer cells grow in approximately spherical 3D aggregates, and exhibit signalling and metabolic profiles more similar to real tumours than is observed in monolayer approaches [17–19].

Like monolayers, spheroids are relatively easy to culture, and growing interest has seen them used for a variety of purposes, including radiobiological application as a means to test fractionation [20–23], as a model for drug delivery [24–28], for investigation of the stem-cell hypothesis [29] and for exploring FDG-PET (Fludeoxyglucose positron emission tomography) dynamics [30] for hypoxia in solid tumours. Crucially, the non homogeneous oxygen distributions in tumour spheroids have been well studied [31–33]. Research to date shows cellular oxygen consumption rate (OCR) has a known influence on the oxygen concentration throughout a spheroid, and directly influences the extent of central anoxia, and the viable rim thickness by known mathematical relationships [32], and thus measuring these aspects allows an experimenter to determine OCR with relative ease compared to other methods [28].

Such methods and the underlying theory are exceptionally important to understanding the factors that influence tumour oxygen distribution, yet all these methods rely on an implicit assumption of perfect sphericity. There is a clear rationale behind this, as symmetry considerations simplify the problem greatly. Yet in experimental conditions, imperfect spheroids are common, frequently growing as extended ellipsoids. When the eccentricity of these shapes is extreme, an experimenter may reasonably choose to discard them from analysis. But this prompts a question; quite how extreme do such deformations have to be before a spherical assumption breaks down? Presumably small departures from sphericity should not impact analysis, whereas highly eccentric ellipsoids could reasonably be presumed to violate the underlying theoretical assumptions. The question of how such eccentricities might skew analysis of spheroids, and how trustworthy results of such analysis might be has not yet been considered in the literature, despite its obvious practical importance.

These questions are as of yet unanswered, and are of paramount importance given the growing adoption of spheroids for cancer research, and their utility in estimating OCR [31,32]. Knowing the acceptable limits of eccentricity for spheroid analysis would be of considerable benefit to experimenters, providing error estimates and limits of reliability. A full analytic expression for “ellipsoidals” (analogous to the spherical case) would also be of substantial benefit, allowing the analysis of eccentric shapes and potentially increasing experimental throughput. In this work, we seek to address these issues by deriving an expression for oxygen diffusion in both prolate and oblate geometries. This is contrasted to the spherical case to determine the limits of validity for experimental data, and the implications of this are discussed. A schematic of this is depicted in Figure 1.

**Fig 1.**
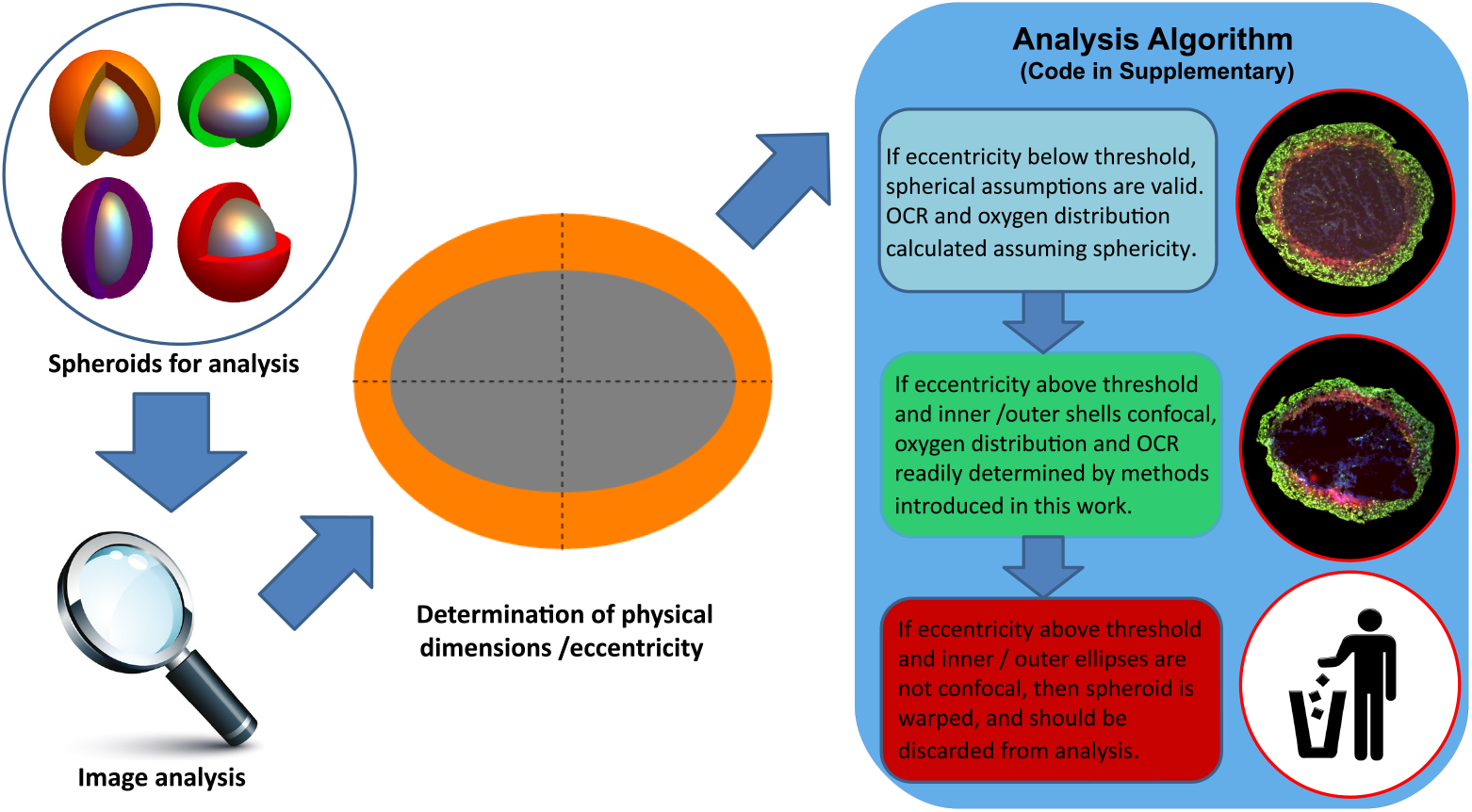
Schematic analysis in this work. Below a calculated threshold for eccentricity, spheroids can be treated as having perfect sphericity without introducing unacceptable error, and their OCR and oxygen distribution established by previously published methods [32]. At greater eccentricities however, a spherical assumption is no longer valid. If the inner and outer sections are concentric ellipses, these can be analyzed by the methods outlined in this work to ascertain OCR and oxygen distribution. If eccentricity is higher than a threshold value, and inner and outer ellipses are not concentric, this suggests the spheroid is severely warped or the section is off the central axis, and should be discarded from analysis. See text for details.

## Spheroids and Ellipsoids

The general equation of an ellipsoid is given by

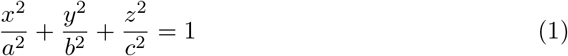

where *a, b* and *c* are the major axes’ lengths. For an ellipsoid with azimuthal symmetry, *a* = *b*. Where all axis are equal (*a* = *b* = *c*), the result is a perfect sphere. To date, this is the only case which has been well-studied from a theoretical standpoint [31–33]. Whilst the mathematical treatments to date have assumed spheroids are perfect spheres, the nomenclature ‘spheroid’ still applies to the more general case, including prolate and oblate spheroids. In this work we broaden the mathematical framework to be applicable to ellipsoids without full spherical symmetry, which are namely

1. Prolate spheroids: In the case where *c > a*, the resulting ellipsoid is an ellipse rotated around its major axis, the line joining its foci. This yields a rugby ball type shape.
2. Oblate spheroids: where *a > c*, an oblate spheroid results, equivalent to an ellipse rotated around its minor axes. The resulting shape is discus-like.

Examples of these ellipsoids are shown in Figure 2. To avoid confusion, we use the term spheroid to refer to an ellipsoidal collection of cells, although we do not limit this to perfect spheres, qualifying with terms ‘prolate’ or ‘oblate’ as appropriate. We use the term ellipsoid to refer to surfaces of iso-concentration of oxygen.

**Fig 2.**
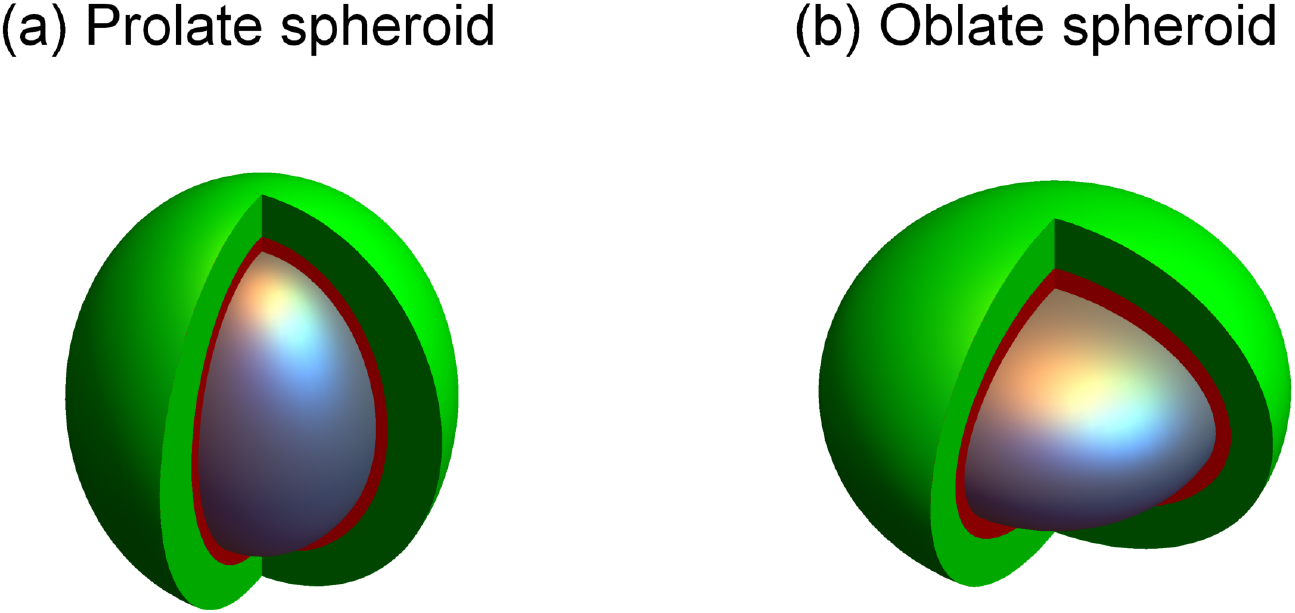
(a) Prolate spheroid (b) Oblate spheroid. The anoxic central core in both cases (*e* = 0.75) is depicted in gray, and the hypoxic extent in red, whilst well oxygenated cells are shown in green. Both ellipsoids have the same volume necrotic core, but their resultant oxygen distributions are slightly different. See text for discussion.

## Model derivation

The full mathematical derivation for oxygen partial pressure in prolate and oblate spheroids is rather involved, and here we shall confine ourselves to stating results with a cursory outline of how they derived. A full mathematical outline is provided in appendix **S1**. Essentially, we are concerned with solving a steady-state reaction-diffusion problem for oxygen field *P* of the form

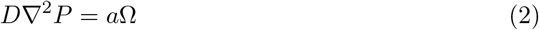

where *D* is the oxygen diffusion constant in water (typically *D* = 2 × 10^−9^ m^2^/s) and *a*Ω is oxygen consumption rate in mmHg/s. This must be solved subject to two crucial boundary conditions, namely that the surface flux and oxygen partial pressure at the anoxic boundary must both be zero. For simple geometries such as perfectly spherical spheroids and cylindrical vessels, symmetry can be exploited to readily yield analytical solutions [33]. In elliptical geometry, the problem is more involved but the basic premise remains the same, and is outlined below. The geometry of the problem is illustrated in Figure 3.

**Fig 3.**
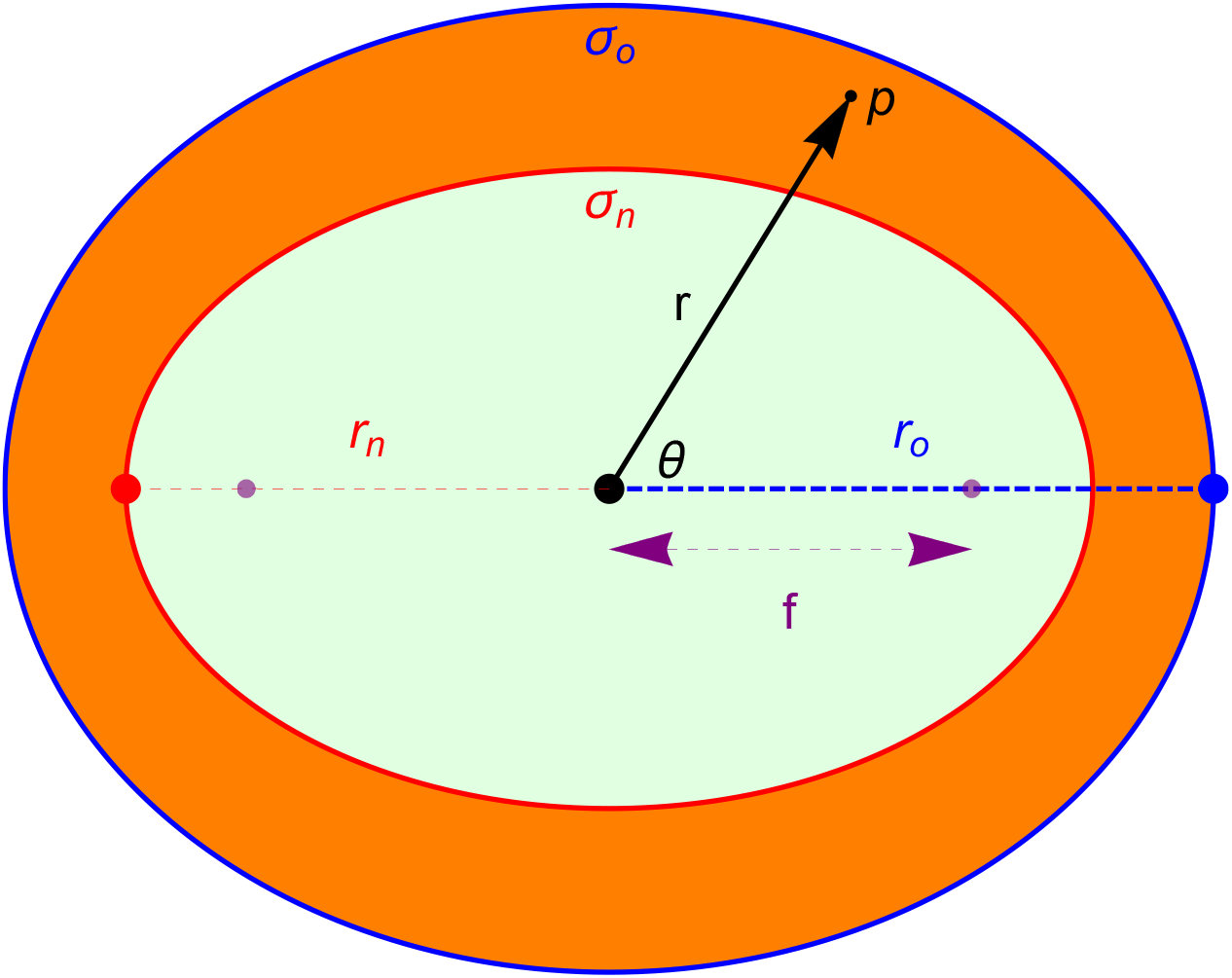
Geometry of a spheroid with outer semi-major axis length *r_o_* and inner semi-major axis length *r_n_*. On the surface *σ_n_*, both partial pressure and oxygen flux are zero. On outer surface *p*(*σ_o_*) = *p_o_*. In spherical co-ordinates, a point *p* is specified by a radial distance from centre *r* and an angle *θ*. The focal length of the inner spheroid is *f*.

### Prolate spheroids

In a prolate spherical geometry, we employ the prolate spherical coordinate system, using a geometrically intuitive definition where curves of constant *σ* are prolate spheroids, whilst curves of constant *τ* correspond to hyperboloids of revolution [34]. This is outlined in detail in supplementary text **S1**. This yields an analytical solution, which can be converted directly into spherical co-ordinates to yield

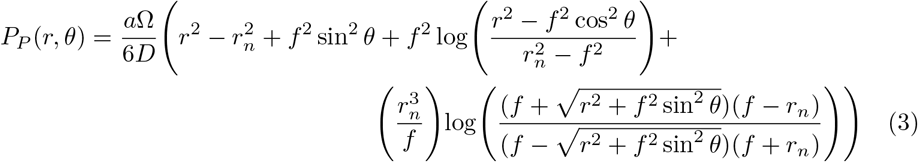

where *f* = *er_n_*, the distance from ellipse centre to foci.

### Oblate spheroids

In oblate spherical geometry, a similar geometrical definition exists [35] and can be solved through similar methods, also outlined in supplementary material **S1**. The full solution in spherical co-ordinates is

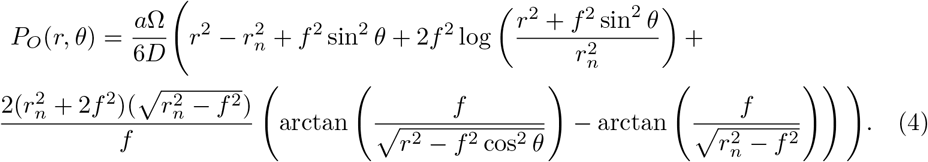

Both the prolate and oblate form can be alternatively cast in terms of *r_o_*, the outer semi-major axis length if preferable. These forms are also given in **S1**.

### Ellipsoidal confocality

Analogous to the perfect spherical case, confocal elliptical surfaces in a spheroid are at the same oxygen partial pressure. For confocal ellipsoidal shells, focal length is constant, related to the eccentricity *e_c_* and semi-major axis of the shell *r_c_* by

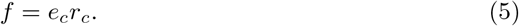

It follows that the innermost (anoxic) and outermost ellipsoidal shells are confocal, thus for a true spheroid *e_o_r_o_* = *e_n_r_n_*. Within the bounds of acceptable experimental error, this relationship can be used to determine whether a given spheroid displaying apparent eccentricity is a ellipsoidal or not. This is important from an experimental perspective, as sectioning can introduce serious distortions in fixed spheroid sections, or can miss the central axis of the spheroid [32]. In these cases, an ostensible ellipsoidal shape might be observed, but may in fact be a sectioning distortion or off-centre cut. Testing for confocality thus determines the underlying reality.

### OCR estimation in spheroids

In the perfectly spherical case, OCR (in mmHg/s) is related to the anxoic radius *r_n_* and outer radius *r_o_* [28,32] by

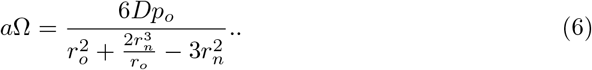

For a prolate tumour spheroid, it is possible to estimate OCR in a manner analogous to the spherical case by re-arranging the equations for *P_P_* to yield

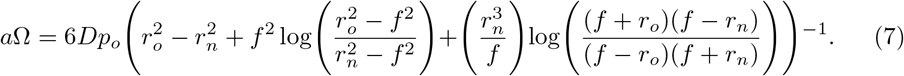

Similarly for oblate spheroids, OCR is given by re-arrangement of *P_O_* to arrive at

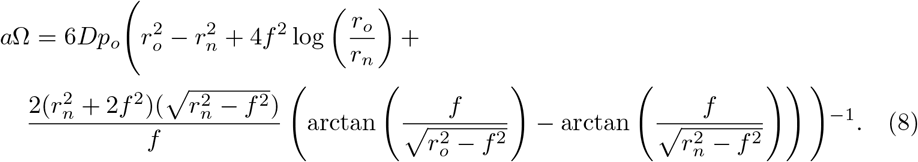

One complication that may arise is that it may be impossible to ascertain whether an ellipsoidal spheroid is prolate or oblate. In that case, one can produce a ‘combined’ expression for average OCR by taking the average of equations 3 and 4, re-arranging to arrive at

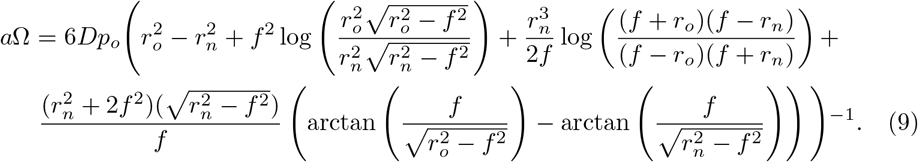

An example of the implementation of these forms including error analysis is included in the code included in supplementary material **S2**.

### Spherical error metrics

It is worthwhile to introduce metrics to quantify how divergent the estimated oxygen-profile in a given ellipsoidal spheroid is from a related perfectly symmetric spheroid. A perfect spheroid has radial symmetry, and thus *P*(*r_o_*) = *p_o_* at all points. Consider related prolate and oblate spheroids both with semi-major axis ro and eccentricity e, nested inside a sphere of radius *r_o_*. We can define the root mean square error (RMSE) by contrasting the expected outer-shell partial pressure p_o_ with what would be measured for spheroids at *P_P_*(*r_o_,θ*) and *P_O_*(*r_o_,θ*) respectively. The RMSE error for prolate and oblate spheroids respectively is

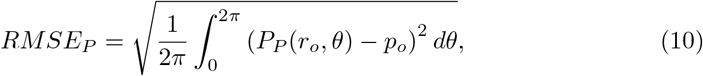

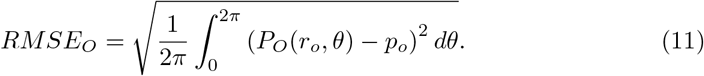

These equations can readily be solved by numerical integration methods, and solutions are demonstrated in supplementary code **S2**. Percentage error is simply *100*(*RMSE/p_o_*), and thus the variation in RMSE with eccentricity can be readily calculated. The other instance when deviation from spherical assumptions must be quantified is in OCR calculation. For example, when the OCR in an spheroid is calculated assuming the perfectly spherical form in equation 6 rather than a more appropriate prolate or oblate form. This might occur when eccentricity is low and the spheroid appears to be entirely symmetric to a first approximation. The distance from the centroid of an spheroid with semi-major axis *r* and focal length *f* to a point on the spheroid at an angle *θ* is given by 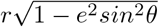, and thus the average value for outer radius *r_o_* and anoxic radius *r_n_* are given by integrating this over a full revolution, yielding

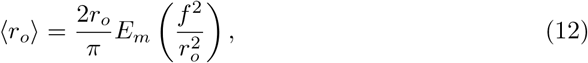

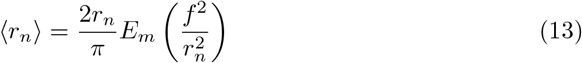

where *E_m_* is the complete elliptic integral of the second kind. Analogous to the discrete standard deviation, the distance function can be integrated over a full rotation 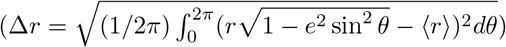 to yield an expression for standard deviation across these spheroids of

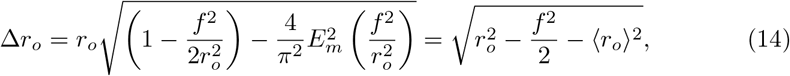

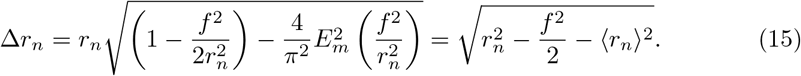

Applying the form in equation 6 for spherical OCR yields a approximation (which is incorrect when *e* > 0) of

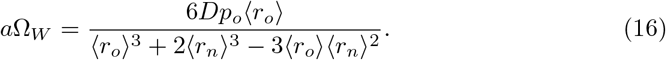

The uncertainty calculation associated with this can be calculated with the variance formula, in this case given by

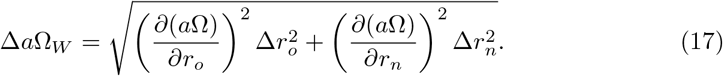

This is analytically tractable, and is given in supplementary material **S1**. An implementation in several code languages is also provided in supplementary **S2**.

## Methods

### Spheroid oxygen profiles

The models derived in this work were used to create oxygen profiles for spheroids of both prolate and oblate classes, which were contrasted to conventional perfectly spherical profiles.

### Quantifying differences between prolate and oblate cases

Prolate and oblate forms have some mathematical differences as can be seen inspection of equations 3 and 4. Whether this difference is experimentally significant is an important question; from a single projection of a tumour spheroid it might be impossible to ascertain whether an experimenter is dealing with a prolate or oblate case. As an experimentalist might not be able to determine whether a given spheroid is prolate or oblate from a single section, quantifying differences in measured OCR under each assumption is an important goal of this work. This was simulated by producing prolate and oblate spheroids with properties as outlined in table 1, and observing the differences in their profiles. In addition, OCR estimates under the ‘wrong’ assumptions were also calculated and inspected. Specifically, the ‘wrong’ assumption occurs when one either applies oblate equations for a prolate spheroid or vice versa. OCR was also calculated with the combined assumption (equation 9) which can also be used when underlying form is unknown. These spheroids were produced with physical properties as per Table 1, with OCR given by equations 6 – 9.

**Table 1.** Simulation parameters

| Parameter | Simulation value |
| --- | --- |
| Semi-major axis | 500 $\mu\text{m}$ |
| Oxygen consumption rate ($a$) | 20 mmHg/s |
| External partial pressure ($p_o$) | 100 mmHg |

### Comparisons with the spherical case

As OCR from spheroids is estimated assuming spherical symmetry, a major aspect of this work was quantifying precisely how close to perfect sphericity spheroids must be so that such an assumption holds, and how departures from sphericity impact estimates of OCR and oxygen profiles. To study this, spheroids with known OCR and varying eccentricity were simulated, and analysed with equations 10–17.

### Experimental proof of concept

To date, non-spherical tumour spheroids have been somewhat neglected, frequently discarded from analysis due to their inherent uncertainty. It is thus difficult to find non-spherical tumour spheroid data. A potential example was taken from a previously analyzed set of sectioned DLD-1 tumour spheroids [32], dual-stained with proliferation marker Ki-67 and EF5. Spheroids from this set were experimentally determined to have an OCR of 22.10 ± 4.24 mmHg/s. The sample spheroid was excluded from prior analysis because of its high eccentricity (external eccentricity *e* ≈ 0.66). This was then analyzed using methods outlined in this work as a proof of concept to determine OCR, contrasting it to known values and spherical estimates.

## Results

### Oxygen distributions in spheroids

Eccentric spheroids were simulated with properties shown in Table 1. Unlike the spherical case, oxygen profiles here are not radially symmetric, so profiles were plotted along both the semi-major and semi-minor axis for clarity. Examples of these profiles are depicted in Figure 4. Oxygen gradients through spheroids are simulated in Figure 5, for both prolate and oblate cases with increasing eccentricity.

**Fig 4.**
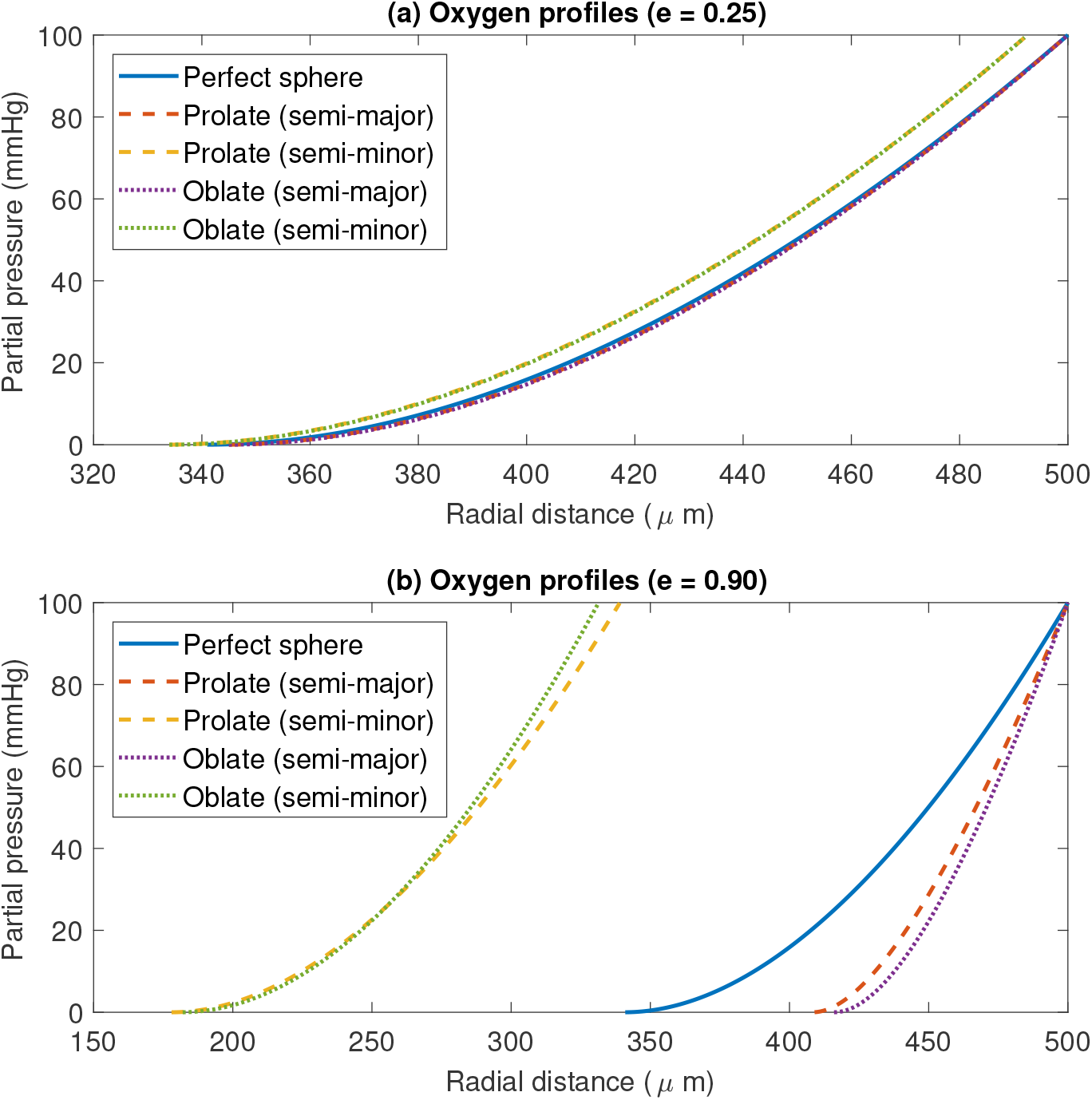
Oxygen profiles for both prolate and oblate spheroids along the semi-major and semi-minor axis for (a) Spheroid with inner eccentricity *e* = 0.25. There is relatively littler difference between prolate and oblate cases and profiles largely overlap, being close to spherical case (b) Spheroid with inner eccentricity *e* = 0.90. Small differences between prolate and oblate cases can be readily seen, and very large divergence from perfectly spherical case is apparent.

**Fig 5.**
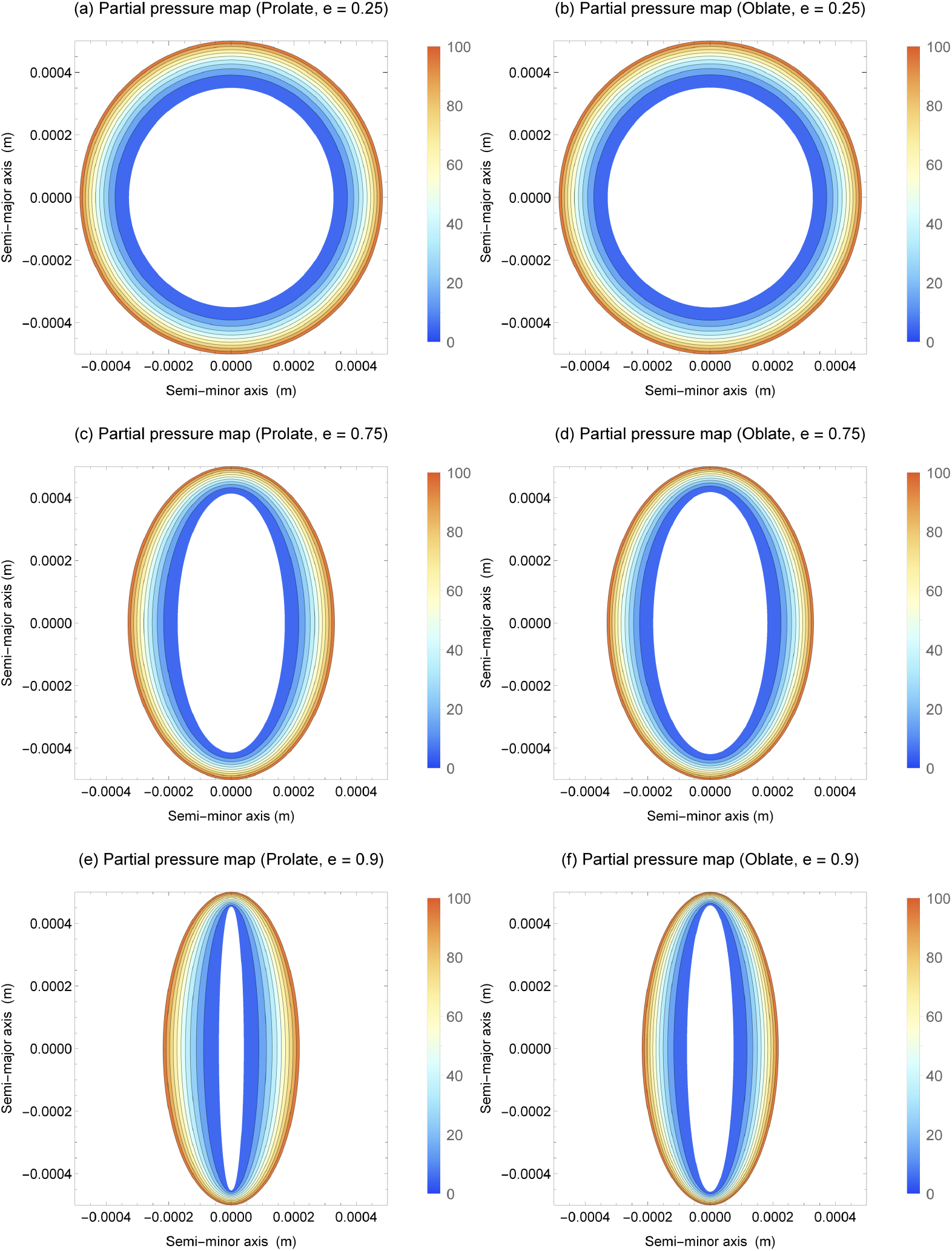
(a) - (f) - Contour oxygen maps for prolate and oblate spheroids with increasing eccentricity with semi-major axis 500*μ*m and OCR of 20 mmHg/s. Prolate and oblate have broadly similar oxygen maps, with divergence manifesting as *e* → 1

### Quantification of differences in prolate and oblate spheroids

From 2D sectioning or imagining alone, it can be experimentally difficult to ascertain whether a given spheroid is either prolate or oblate. It is thus important to quantify differences between the ellipsoids. Figures 4 and 5 suggest that prolate and oblate spheroids have broadly similar oxygen profiles until eccentricity approaches unity. Figure 6 depicts internal anoxic radii with eccentricity. These differ only slightly, typically ≤ 1% for both major and minor axis for 0 < *e* < 0.9. More interesting perhaps is the variation in OCR estimate with eccentricity under the ‘wrong’ assumption (namely assuming oblate form when actual entity is prolate or vice versa) shown in Figure 7. This suggests strongly that OCR estimates arrived at under the ‘wrong’ assumption are still accurate up until high eccentricity. Even at very high eccentricity, an average value of both incorrect OCRs was extremely close to true OCR. This suggests that incorrectly specifying the type of spheroid should not greatly impact OCR estimates. For improved accuracy, employing the combined estimate in equation 9 yields even smaller errors, suggesting this could readily be used by experimentalists without introducing major error even when the underlying form is unknown.

**Fig 6.**
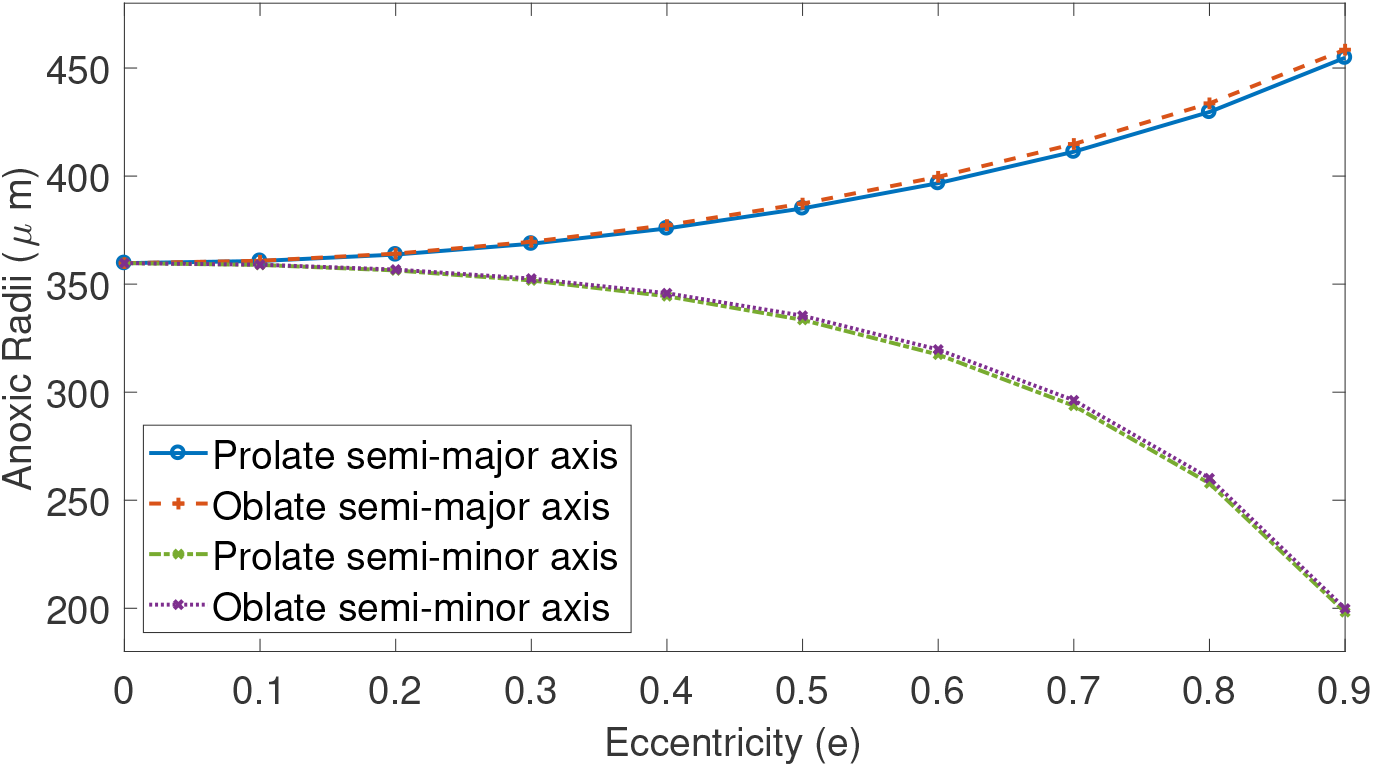
Variation of anoxic radii with eccentricity of outer shell from equations 3 and 4 for spheroid with properties as per Table 2. Differences between radii in prolate and oblate case are generally very small up to high eccentricity.

**Fig 7.**
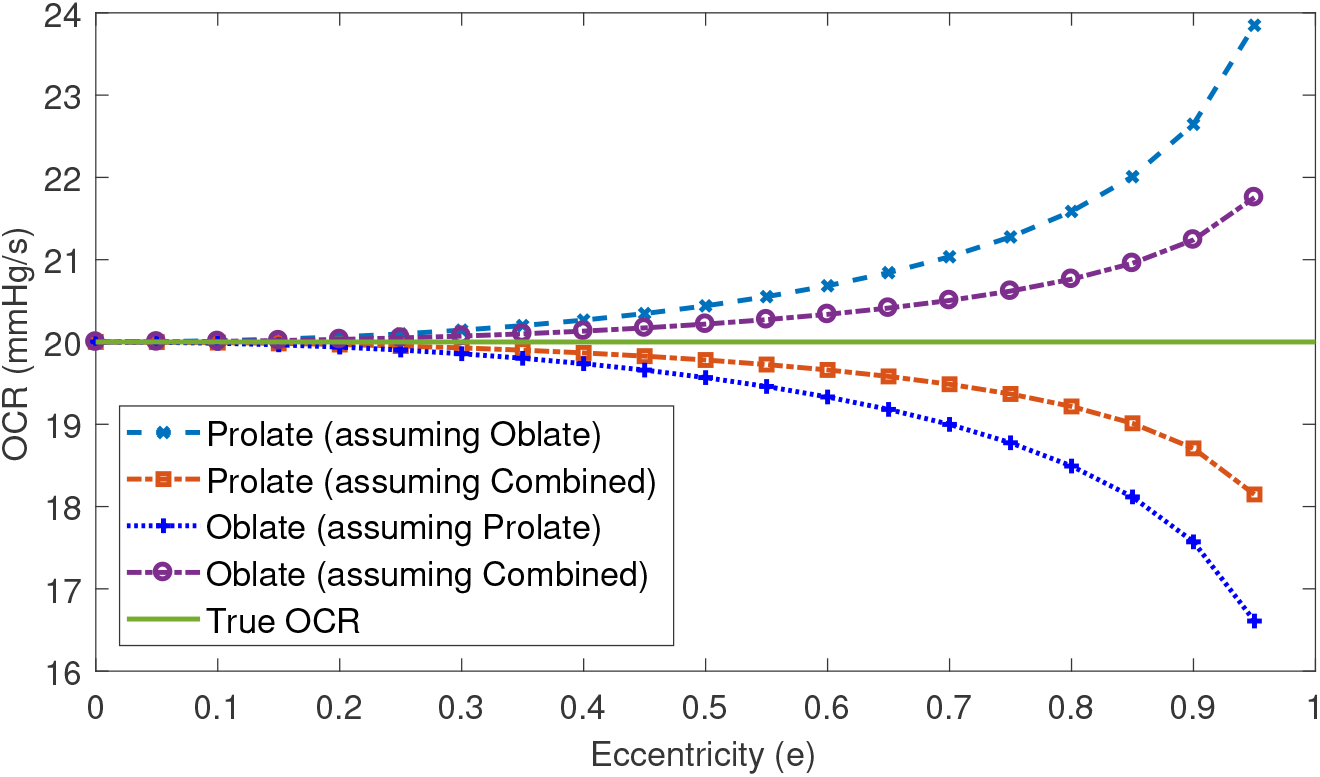
OCR estimates under ‘wrong’ assumptions for increasing inner eccentricity, calculated from equations 7 – 9. Combined OCR estimate yields smaller errors in all cases. True OCR is 20 mmHg/s.

### Comparisons with spherical case

Table 2 depicts the impact of assuming perfect sphericity on derived OCR estimates for properties in Table 1. At low *e* (typically *e* ≤ 0.3), treating spheroids as perfect spheres yields acceptable accuracy for both OCR estimates. However, as *e* → 1, the reliability of OCR estimates rapidly break-down, and errors become increasingly large and unreliable.

**Table 2.** RMSE error and OCR estimates assuming sphericity for spheroids of varying eccentricity *e*

| $e$ | RMSE<br>(Prolate) | RMSE<br>(Oblate) | Prolate OCR<br>(mmHg/s) | Oblate OCR<br>(mmHg/s) |
| --- | --- | --- | --- | --- |
| 0 | 0 % | 0 % | $20.00 \pm 0.00$ | $20.00 \pm 0.00$ |
| 0.1 | 0.82 % | 0.82 % | $20.13 \pm 0.37$ | $20.14 \pm 0.37$ |
| 0.2 | 3.32 % | 3.33 % | $20.54 \pm 1.53$ | $20.60 \pm 1.54$ |
| 0.3 | 7.77 % | 7.78 % | $21.27 \pm 3.73$ | $21.42 \pm 3.78$ |
| 0.4 | 14.57 % | 14.60 % | $22.44 \pm 7.47$ | $22.75 \pm 7.68$ |
| 0.5 | 24.45 % | 24.55 % | $24.26 \pm 13.91$ | $24.84 \pm 14.56$ |
| 0.6 | 38.67 % | 38.94 % | $27.16 \pm 25.76$ | $28.25 \pm 27.76$ |
| 0.7 | 59.51 % | 60.17 % | $32.25 \pm 51.07$ | $34.41 \pm 57.82$ |
| 0.8 | 91.53 % | 93.12 % | $43.16 \pm 124.77$ | $48.58 \pm 156.72$ |
| 0.9 | 146.17 % | 150.33 % | $85.56 \pm 660.92$ | $120.52 \pm 1291.81$ |

### Experimental proof of concept

For this work, a simple image analysis algorithm was written for the spheroid image, which found the ellipsoid centre and cast best fit ellipses from this position. The analysis algorithm was broadly similar to previously described methods [32], yielding an estimates of *e* ≈ 0.66, *r_o_* = 488.5*μ*m and *r_n_* = 400.15 *μ*mm. This analysis suggested the best-fit inner and outer ellipses were approximately confocal to within experimental error as required by confocality condition in equation 5 (*e_o_r_o_* ≈ *e_n_r_n_* to within an error of 1.13%). Uncertainty on the lengths of *r_o_ r_n_* were taken from this to be 5.53*μ*m and 4.52*μ*m respectively. Results of this analysis is depicted in Figure 8 and Table 3, suggest values in agreement to those previously measured when considered as an prolate / oblate spheroid, and unrealistic values if presumed perfectly spherical.

**Fig 8.**
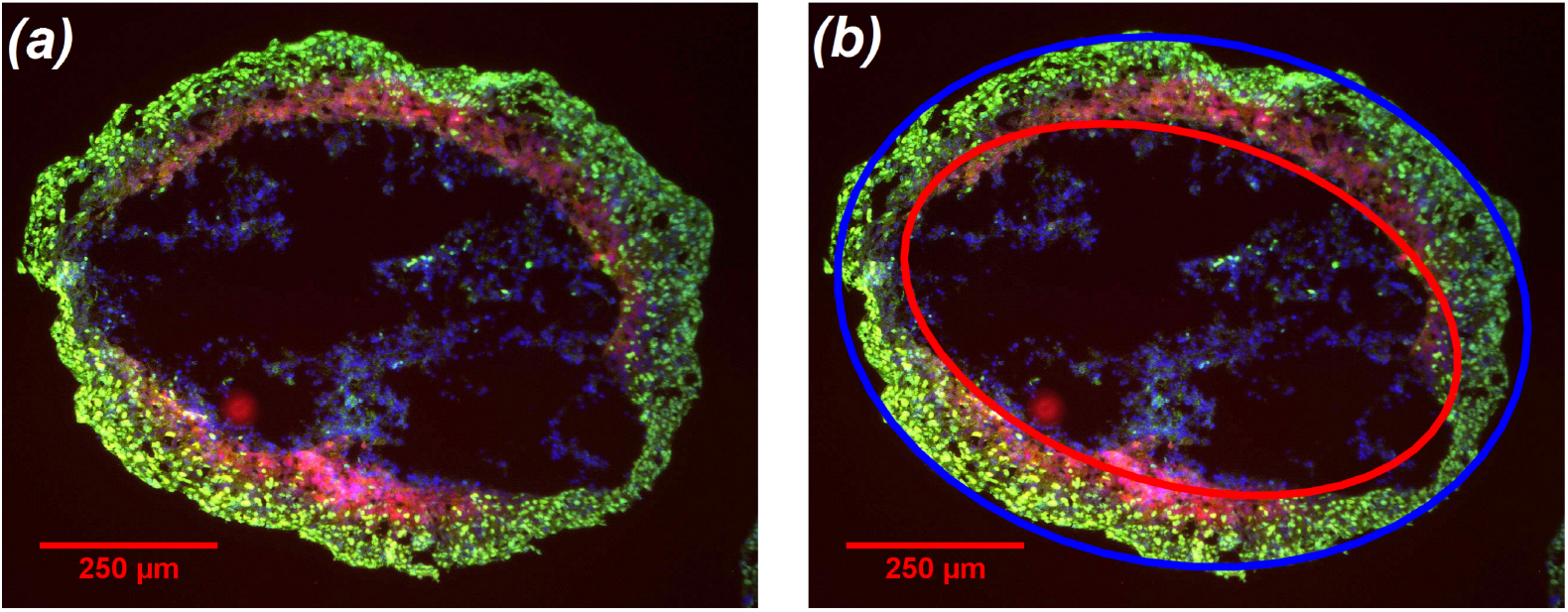
(a) Eccentric DLD-1 spheroid (contrast enhanced for clarity) (b) Demonstration of analysis in with a spheroid image-analysis algorithm detecting best-fit ellipses (blue ellipse best-fit to outer boundary, red to inner). Calculated focal lengths from both are checked for confocality from equation 5 and OCR estimated. See text for details.

**Table 3.** OCR estimates for sample spheroid

| Assumption | Estimated OCR (mmHg/s) |
| --- | --- |
| Previously measured for DLD-1 cell-line | $22.10 \pm 4.24$ |
| Assumption of perfect sphericity (true spheroid) | $42.74 \pm 51.98$ |
| Assuming prolate tumour spheroid | $29.25 \pm 4.25$ |
| Assuming oblate tumour spheroid | $27.27 \pm 4.11$ |

## Discussion

Analysis of spheroids to date tends to pivot on the presumption of sphericity, as symmetry arguments reduce the complexity required. However, real spheroids tend to depart from perfect sphericity to varying extents. In this work, we provide a metric for determining how reliable the simpler spherical assumption will be as eccentricity increases, as outlined in Table 2. The methods outlined in this work can be employed to generate reliable estimates of OCR and oxygen distribution. Another major benefit of this work is that it allows an experimenter to determine OCR even in non-spherical cases when high eccentricity might otherwise render the spheroids in question ill-suited for analysis. Provided the inner and outer ellipses are suitably confocal, the analysis outlined in this work can be employed, and thus should help increase experimental throughput.

As depicted in the Table 2, presuming sphericity with eccentric sections yields acceptable accuracy when eccentricity is small, but rapidly begins to produce completely unrealistic results for OCR and massive uncertainty. When analysed as either prolate or oblate spheroids however, OCR estimates are in agreement with the previously measured values. This suggests strongly that for eccentricity greater than approximately e = 0.3, spherical assumptions for OCR cease to be appropriate and ellipsoidal analysis must be employed. While data for this is currently sparse, we were able to demonstrate the principle on the highly eccentric spheroid illustrated in Figure 8, determined by image analysis to have *f* = 318.89 ± 2.57μm. When analysed as a spheroid, OCR estimates were completely unrealistic with huge uncertainty, as seen in Table 3. However, when considered as either a prolate or oblate spheroid, OCR measurements were within previously measured values. This is promising, but as only a single data point is available, this should be interpreted solely as a proof of concept.

It’s worth noting that from single section images or microscopy, there is no obvious way to ascertain whether a spheroid is prolate or oblate. As the mathematical forms for these are slightly different, this adds an extra uncertainty and prompts the question about which form is preferable to employ. The analysis in this work (Figures 6 and 7) indicate that even if one incorrectly assumes the wrong form, OCR estimates are still very good, with only minimal errors introduced. This holds with only negligible errors until very high eccentricity. The combined OCR form in equation 9 yields only negligible error even when the underlying form is unknown. Thus an experimenter should opt to use this form for OCR estimation when they have no other information on whether the specimen is prolate or oblate, as this does not introduce large errors even at high eccentricity.

While modelling of elliptical oxygen diffusion has the potential to greatly extend experimental throughput, there are a number of scenarios where an ostensible eccentric spheroid might not be what it appears. For fixed and sectioned spheroids, the act of sectioning itself can be enough to induce substantial deformations, stretching it along a particular axis. Ostensibly, the resultant shape might appear ellipsoidal, but is in reality a warped spheroid, and cannot be reliably analysed with the methods outlined. Such an example is show in figure 9, for a spherical spheroid sheared along an axis. From the mathematics established in this work, we can distinguish between true spheroids and warped spheroids - if the inner and outer ellipses are not confocal (*e_o_r_o_ = e_n_r_n_*), then the shape is a warped spheroid, and should be discounted from analysis, as per Figure 1. Crucially, Figure 9 demonstrates that warped spheroids can only satisfy the ellipsoidal confocality condition under two circumstances; either when its eccentricity is 0 (a perfect sphere), or the non-physical situation where *r_o_ = r_n_*. Thus a sheared spheroid in one direction will never come close to satisfying the ellipsoidal confocality condition. In practice, all experimental work comes with inherent uncertainty, so *e_o_r_o_ ≈ e_n_r_n_* within the bounds of image analysis uncertainty is sufficient to determine whether a spheroid can be treated as an ellipsoidal case.

**Fig 9.**
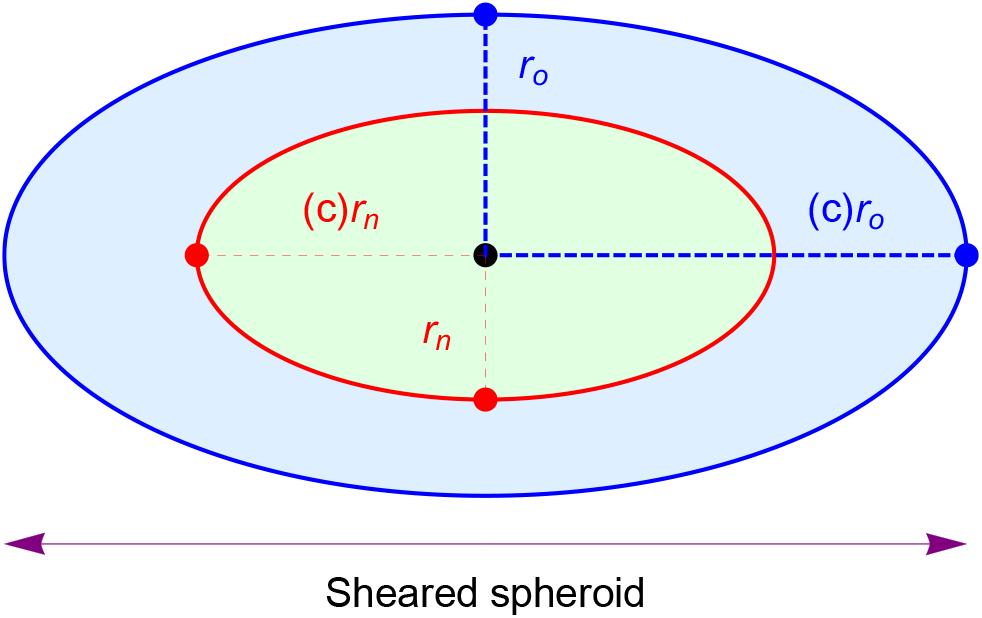
The eccentricity of a sheared spheroid is the same for both anoxic and outer ellipsoids 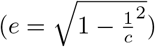 and thus not confocal as *f* is not constant. Sheared spheroids are not true spheroids, as confocality condition is not met.

There is a more subtle issue with sectioned spheroids, which becomes even more crucial with sectioned eccentric spheroids. Analysis relies on a section through the central axis of the spheroid. In the perfectly spherical case, if the section is off-centre, the net result will be two concentric circles but with a misleading ratio, rendering any OCR calculation derived from this suspect. By contrast, any plane through an ellipsoid produces an ellipse, but if these cuts are off the central axis, then the inner and outer ellipses will not longer have a common centre, and will not be confocal. In this regard, determining an off-centre ellipsoid section is relatively straight-forward. A proof of this is provided in supplementary **S1**.

The theoretical analysis outlined here presents biological investigators with new methods for extending spheroid analysis, and means to interpret data which departs from sphericity. It also establishes uncertainty bounds on existing spherical analysis techniques, and methods for determining OCR and oxygen distribution in tumour ellipsoids. The findings of this work will increase experimental confidence with tumour spheroids, and have the potential to substantially increase experimental throughput, improving our insights on everything from tumour hypoxia to drug delivery. Such an approach is imperative if we are to fully exploit this unique experimental tool, and ultimately marshal the new insights obtained towards better cancer treatment and diagnostics.

## Supporting information

**S1** Appendix. Mathematical appendix and full derivations.

**S2** Appendix. Sample code for ellipsoid oxygen distribution and OCR calculation (Mathematica, MATLAB / Octave, Excel spreadsheet)

## Author contributions

DRG and FJC conceived the theory outlined in this work, DRG derived the model, performed analysis, and wrote the manuscript. Both authors reviewed the manuscript and authored supplementary texts.

## Data availability

All data is available with the paper, and sample implementations in MATLAB, OCTAVE, Mathematica, and Excel spreadsheet are included in supplementary material **S2**.

